# Toward More General Embeddings for Protein Design: Harnessing Joint Representations of Sequence and Structure

**DOI:** 10.1101/2021.09.01.458592

**Authors:** Sanaa Mansoor, Minkyung Baek, Umesh Madan, Eric Horvitz

**Affiliations:** Department of Biochemistry and Institute for Protein Design, University of Washington, Seattle, WA 98195, USA; Microsoft, Redmond, WA 98052, USA

## Abstract

Protein embeddings learned from aligned sequences have been leveraged in a wide array of tasks in protein understanding and engineering. The sequence embeddings are generated through semi-supervised training on millions of sequences with deep neural models defined with hundreds of millions of parameters, and they continue to increase in performance on target tasks with increasing complexity. We report a more data-efficient approach to encode protein information through joint training on protein sequence and structure in a semi-supervised manner. We show that the method is able to encode both types of information to form a rich embedding space which can be used for downstream prediction tasks. We show that the incorporation of rich structural information into the context under consideration boosts the performance of the model by predicting the effects of single-mutations. We attribute increases in accuracy to the value of leveraging proximity within the enriched representation to identify sequentially and spatially close residues that would be affected by the mutation, using experimentally validated or predicted structures.

## 1 Introduction

Proteins are molecular machines that perform nearly all of the critical functions in living organisms. They consist of linear chains of amino acids that spontaneously fold into unique three-dimensional structures which carry out biochemical functions. The sequence to structure to function relationship is integral in understanding how to design proteins for specific functions. Most methods that seek to understand the complex sequence to structure to function relationship take either a first principles approach with structural simulations [1] or leverage sequence embeddings trained through an adaptation of semi-supervised machine learning methods used to construct large-scale neural models developed for natural language processing (NLP) tasks [2, 3, 4, 5].

Sequence embeddings created via semi-supervised training have demonstrated strong performance over a broad range of biologically relevant downstream tasks, particularly in the realm of protein engineering [6, 7]. Through the semi-supervised training objective, these models capture long-range dependencies between unrelated families of proteins. Protein structure is more informative for predicting function than sequence [8]. However, the relatively small number of structurally-validated proteins, as compared to the expansive set of sequences available, motivates the focus of attention to date on protein sequences for protein function prediction tasks. Language models centered on semi-supervised training on sequences continue to grow and become increasingly accurate with larger architectures, more compute time, and more data. We have explored an alternate approach, promising greater data-efficiency via the explicit introduction of structural information. We view this as providing models with strong biological priors.

We present a deep learning framework that explicitly encodes structural information into pre-trained sequence embeddings to form a more informative and generalizable embedding of proteins. Use of the approach led to an improvement in masked sequence and structure recovery. Additionally, we showed that the joint training with sequence and structure information improved predictions of the effect of single mutations on thermal stability. With recent considerable improvement in protein structure prediction techniques [9, 10], this multi-task trained model is able to accurately predict protein properties from structural models, getting around the need of experimentally validated structures.

## 2 Results

### 2.1 Masked Structure and Sequence Recovery

We evaluated the performance of the model for recovering the masked structure on the validation set. The same masking scheme was employed, where continuous regions of 5 residues were masked to make up a total of 15% masked residues. Figure 2.A shows that all the masked validation points were corrected when passed through the model. On average, the initial validation samples were perturbed by 5 angstroms, and post-correction, the samples had an RMSD of about 2 angstroms from their respective native protein. To evaluate whether addition of structure leads to an improvement in sequence recovery, we compared our results with the sequence-only baseline (ESM-1b). The masking for this task was adopted from ESM-1b, where random tokens were chosen to make up a total of 15% tokens masked. Figure 2.B shows the accuracy of the predicted masked amino acid for both the joint embedding model and baseline. Due to the added structural context, the model was able to recover the masked sequence tokens to a higher accuracy than the sequence-only baseline, for most validation points. Average baseline sequence recovery was 0.06 (6%), whereas for the joint embedding model, we were able to achieve an accuracy of 0.13 (13%) on the same test set.

### 2.2 Predicting the effect of a single mutation on thermal stability

To predict the effect of a single mutation on thermal stability, we fine-tuned the sequence prediction head (LMHead) for both the sequence-only baseline and the joint embedding model. Since the architecture of the sequence prediction head is the same, this serves as a head-to-head comparison of the value of the enriched information captured in the embedding space and how well equipped it is to predict the effect of a single mutation. The train and test set consisted of mutants from non-overlapping wild-type proteins. Figure 3.A shows the correlation between predicted and measured ΔΔG values for the same test set. The general embedding model has a higher Spearman rank-order correlation coefficient (spearmanr) of its predictions of ΔΔG to the ground truth than the sequence-only baseline, providing evidence that the explicit addition of structure to the embedding space leads to a more informative protein encoding for the prediction of single mutations.

### 2.3 Evaluation of mutant vs wild-type embedding space

To pursue insights about the operation of the model in predicting the effect of a single mutation, we compared the difference in the embedding space of the wild-type to the mutant protein for both the sequence-only baseline and the joint embedding model (Figure 3.B and Supplemental Figure 5). When plotted on the same scale, we can see that the structure-aware embedding shows a higher degree of difference between the wild-type and mutant embedding spaces for PDB 1FXA. The general embedding model also shows sequentially-distant residues being influenced by the mutation. Inspection of the 3D structure of the protein reveals that these distant sites are structurally close. We consider these findings as evidence that the joint embedding is genuinely structure-aware, making it more sensitive to mutation effects than the sequence-only model.

### 2.4 Single mutation effect prediction using predicted models

Computational methods for predicting the effect of mutations can provide great value for protein design. We evaluated the performance of the model on ΔΔG prediction using structural models, rather than experimentally validated structures. We used RoseTTAFold [10] for predicting models of the ProTherm test set for this experiment. On average, the predicted model structures differed about 1.2 angstroms from their respective native structures (Supplemental Figure 4). Figure 3.A (right plot) shows the correlation between predicted and ground truth ΔΔG for predicted models for the test set. This points to the generalizability of the model to withstand perturbations to the input structure and that the model does not require experimentally determined structures to provide high accuracy predictions of the effect of single mutations. The sufficiency of using predicted structures can significantly raise the efficiency of protein design, with uses, for example, in ranking the efficacy of function of candidates and better triaging them for expensive wet-lab experiments.

## 3 Discussion

The exponential increase in protein sequence databases has led to major developments in developing generative and predictive models for protein engineering using deep learning. Validated protein structures, on the other hand, constitute a small fraction of all known biological sequences. Effectively combining structure and sequence information promises to be beneficial for downstream prediction problems where data is limited. With increasing progress in language modeling for proteins, we have seen that these models continue to benefit from more data and computation power. However, the explicit joining of structure and sequence information in embedding models can provide a more data-efficient approach to encoding proteins that can reduce required quantities of training samples and compute power.

The methodology we have taken to develop a more general embedding for proteins combines both structural and evolutionary information in a semi-supervised manner using a SE(3)-equivariant Transformer. Since the model was trained with a multi-task loss on both the masked sequence and structure prediction from the embedding space, equal weight was given to recovering the sequence and structure. Both the masked sequence tokens and masked structure coordinates were recovered accurately from the 128 dimensional embedding space.

The structure-aware embedding trained was able to capture complex relationships between amino-acids in both sequential and structural space. Our model was more sensitive to single-point mutations in the wild-type protein than the sequence-only baseline, suggesting the value of building richer encoding that contains both protein sequence and structure information. We were able to predict the effect that a point mutation would have on thermal stability of a protein to a higher accuracy than the sequence-only baseline, which provides evidence of the value of adding structural information. Also, when evaluating the difference in the embedding space of the mutant versus the wild-type protein, the general embedding model was able to identify sequentially-far but spatially-close residues that would be affected by the point mutation. Finally, since determining protein structures is expensive and time-consuming, we evaluated the single-mutant effect prediction accuracy of our model using structure models predicted by RoseTTAFold. The mutant effect prediction accuracy remained high even though the structural models were approximate. This points to the generalizability of the method developed and its viable use case for protein design challenges using design models.

We see multiple directions for improvement of the joint use of sequence and structure information for both masked token recovery and downstream prediction tasks. Through multiple iterations with either shared or independent weights, incremental corrections to the structure coordinates would help with masked recovery. Increasing the model size (currently 16 million parameters) through increasing the size of the embedding space would enable the model to store more information from both sequence and structure. In another direction, a promising path to achieving more accurate structure recovery is to revise the approach to predicting corrections for each atom. We predicted corrections for atoms (Ca, C, N, Cb) independently. In an alternate approach, we can treat atom orientation and displacement as interdependent, where all atom corrections are relative to the Ca correction. We see numerous prediction tasks as benefiting from the construction and leveraging of the joint embeddings. We are exploring the use of the methodology and extensions for predicting the probability of a mutation being deleterious using Deep Mutational Scanning (DMS) data.

Overall, this method points to the viability of encoding sequence and structure to form richer and more informative embeddings of proteins. We hope that the model provides inspiration for future development of more general protein embedding models.

## 4 Methods

### 4.1 Input sequence embedding and structure information

We used the trRosetta2 [11] training and validation datasets for our model. The masked sequence input was fed into the ESM-1b model [5], loaded with pre-trained weights. By doing so, we are enabling the generation of already learned sequence representations to be used as input for our model. We used the sequence embedding from the last layer (1D feature) along with the attention maps from each of the 33 layers (2D features) from ESM-1b. For structure, we only considered N, Ca, C, and Cb atoms (virtual Cb atoms for GLY). Proteins longer than 128 residues were cropped to fit the memory of a single NVIDIA V100 (16GB).

### 4.2 Training Objective and Details

We employed a semi-supervised training task, where we masked out random continuous regions of 5 residues to make up a 15% total mask of the input protein. Same regions of the sequence and structure were masked. For masking, we used a special mask token for sequence, and perturbed the structure coordinates by at least 5 angstroms, to make the masked structure sufficiently different from native, with additional noise from a standard normal distribution. The sequence tokens were recovered by minimizing cross entropy loss between the model’s predictions of the masked amino acid and the true amino acid. The structural loss was defined as the batch average of the mean squared error over the model’s predictions of the masked atom coordinates and the true coordinates over the 4 types of atoms considered. The total loss was an addition of a sequence loss and a structural loss weighted equally. The model was optimized using Adam (*β*1 = 0.9, *β*2 = 0.999) with learning rate 1e−3, with an effective batch size of 128.

### 4.3 Model Architecture

We used a SE(3)-equivariant transformer [12], a graph neural network, for processing the masked sequence and structure features (Figure 1). The 1D (sequence embedding) and 2D (attention maps) ESM-1b features were linearly projected down to 32 dimensions for noise reduction. For constructing the graph, each node represented a residue of the masked structure where we considered a neighborhood of the 16 closest neighbors based on perturbed Ca distances. Node features included the ESM-1b sequence embedding for each residue and the displacement vector of each atom to Ca (NCa, CCa, CbCa). Edge features included the ESM-1b 2D attention map and the inter-residue orientations [13] (Supplemental Figure 6). The edge connecting residues 1 and 2 would contain the attention map values for the two residues and the inter-residue orientations represented by 3 dihedral (*ψ, θ*12, *θ*21) and 2 planar angles (*ϕ*12, *ϕ*21). This graph was fed into a 5-layered SE(3)-equivariant transformer, the output of which was a 128 dimensional hidden embedding of the input features. The embedding space was fed separately into a masked sequence prediction head (LMHead) and another SE(3)-equivariant transformer for masked coordinate prediction (SE3-Structure Recovery or SE3-SR) (Figure 1). The architecture of the masked sequence prediction head is adopted from the LMHead of ESM-1b model [5], a 2 dense layered network with layer normalization. For the SE3-SR top model, we used a 3-layered SE(3)-transformer where the node features included the jointly trained embedding from the first SE(3)-transformer (128D), along with the displacement vector of each atom to Ca. The edge features included the inter-residue orientations only.

**Figure 1:**
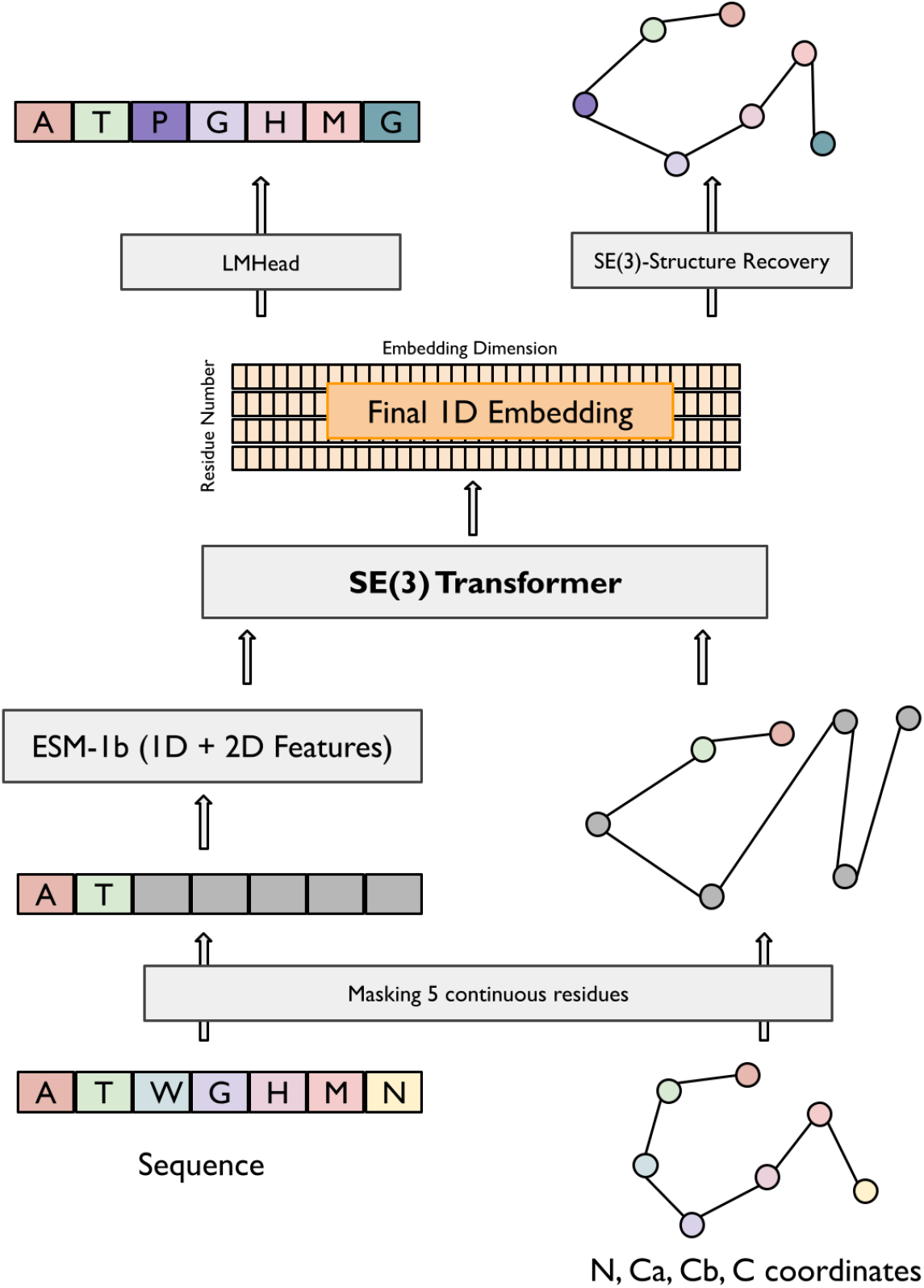
Model architecture for building and leveraging protein embeddings that make joint use of sequence and structure information. Same sequence and structure regions are first masked. Both 1D and 2D features are generated from passing the masked sequence through the pre-trained ESM-1b model. Masked structure and sequence representation are passed as input to a SE(3) transformer, which is trained to output a 128 dimensional embedding space. The embedding space is used to predict the masked sequence and structure regions.

**Figure 2:**
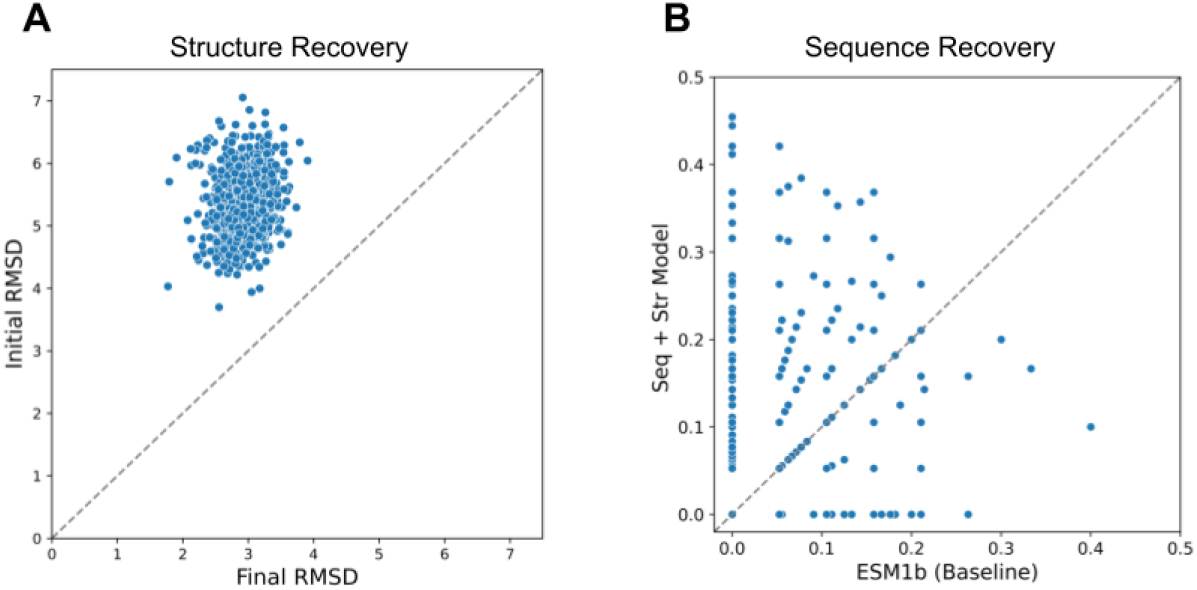
Structure and sequence recovery. (A) Scatterplot of RMSD of masked (perturbed) input structures and RMSD of corresponding corrected structures. (B) Scatterplot comparing sequence recovery of trRosetta validation samples from general embedding model and sequence-only baseline (ESM-1b).

**Figure 3:**
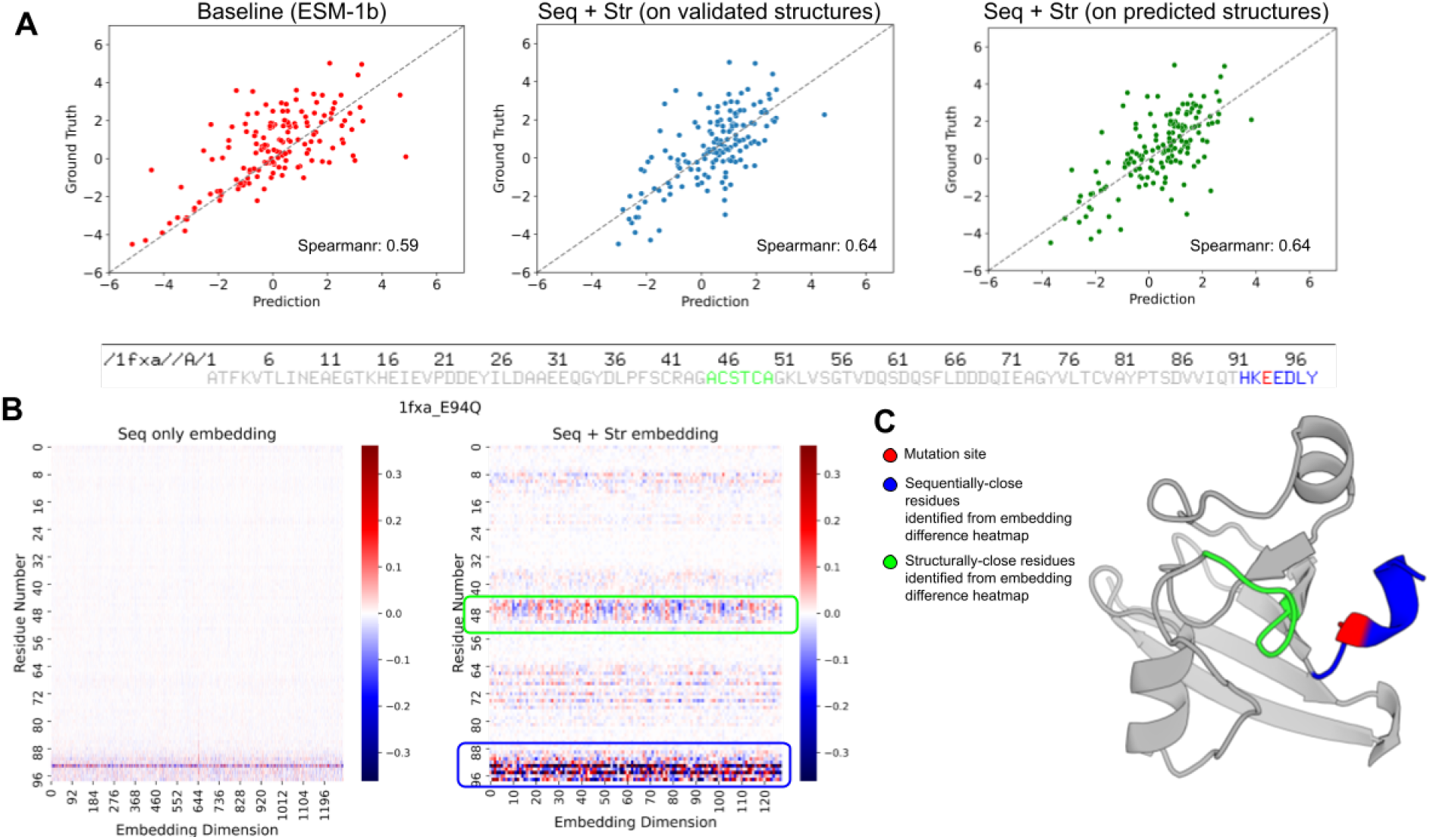
Accuracy of prediction of ΔΔG of single mutants and analysis of PDB 1FXA, with single mutation of residue position 94 (E → Q). (A) Scatterplots showing correlation of predicted ΔΔG to experimental values (ground truth) using sequence-only (ESM-1b), joint sequence and structure embedding model using experimentally validated structures and joint sequence and structure embedding model using predicted structural models from RoseTTAFold. (B) Heatmaps showing the difference in mutant and wild-type embeddings generated for single mutation on residue position 94 for PDB 1FXA, using sequence-only and joint sequence and structure embedding models, with regions having the highest difference in embedding values highlighted in blue and green. (C) 3-D structure of PDB 1FXA with the mutation site colored in red. The most-affected regions identified from the embedding space difference between wild-type and mutant are highlighted using the same color schemes.

### 4.4 Fine-tuning for single-mutant effect prediction

Following semi-supervised training of the model, we fine-tuned our model for the task of predicting the effect of single-mutations on thermal stability (ΔΔG values). A subset of the ProTherm dataset [14] was used which consisted of 1042 mutants from 126 wild-type proteins. Training and test sets consisted of non-overlapping wild-types and their mutants. The training and test set were split 80:20. The fine-tuning task was adopted from the published ESM-1b model, where the mutant position was masked and the predicted value is the difference between the log probabilities of the mutant amino acid and the wild-type amino acid. Only the sequence prediction top model (LMHead) from the embedding space was re-trained for both the joint embedding model and the baseline (ESM-1b). As mentioned previously, the sequence prediction head for both the joint embedding model and the sequence-only baseline has the same architecture.

## 5 Acknowledgments

We would like to thank David Baker, Kevin Yang, Justas Dauparas, Hahnbeom Park, Ivan Anishchanko, Hugh Haddox and Paul Koch for helpful comments and datasets. This work was supported by Microsoft (S.M, M.B, U.M, E.H) and Washington Research Foundation (M.B).

## 6 Supplemental Information

**Supplementary Figure 4:**
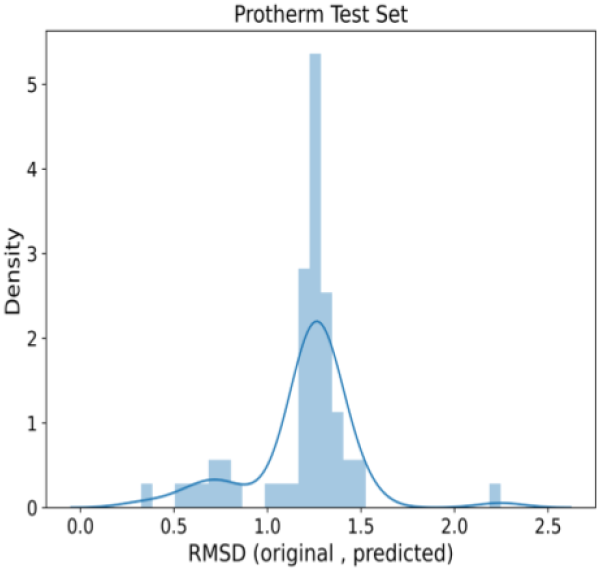
RMSD of predictions on Protherm test set. Distribution of RMSD of experimentally validated structures to their respective RoseTTAFold structure prediction models.

**Supplementary Figure 5:**
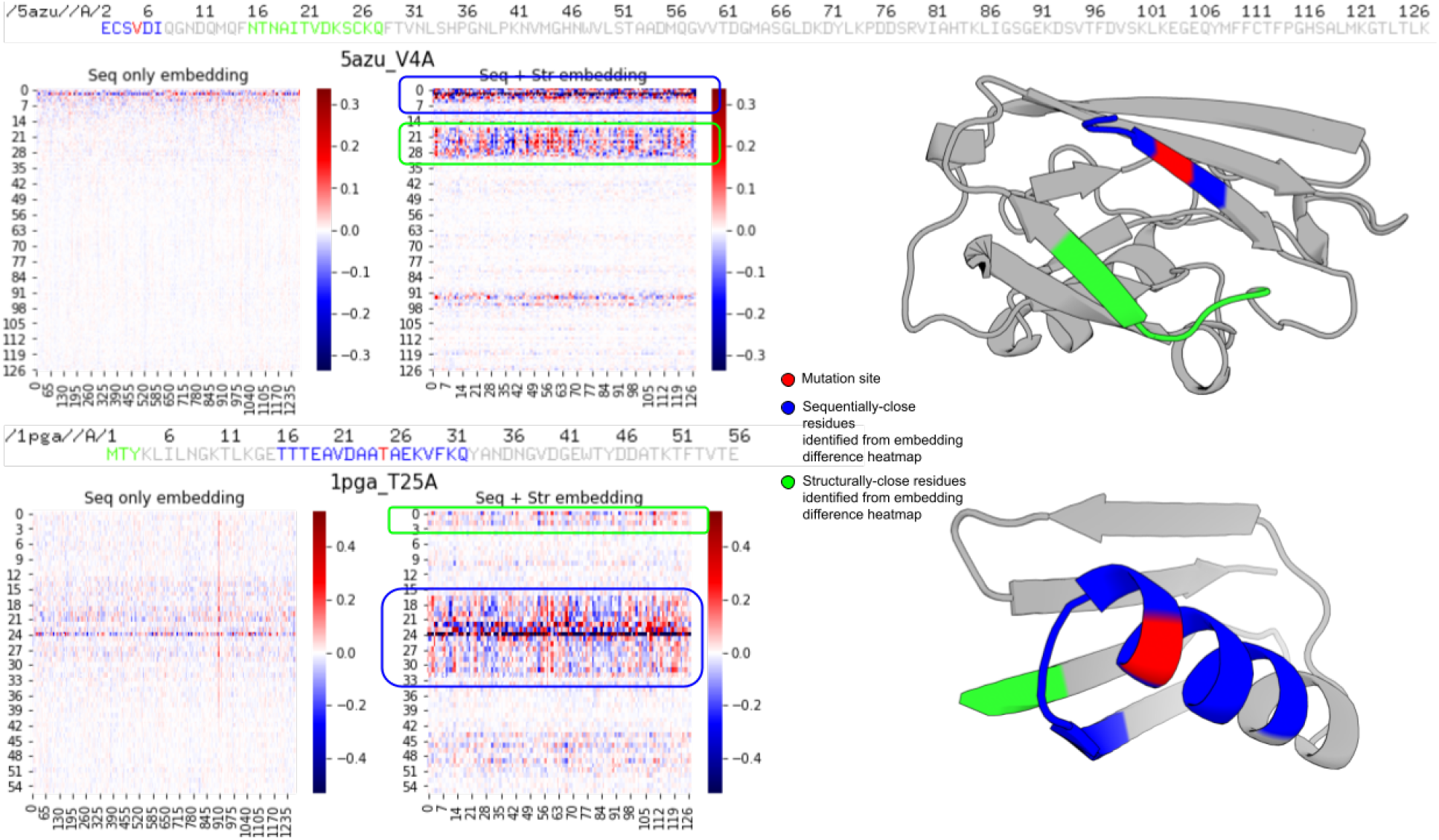
Additional single mutation analysis of PDBs 5AZU and 1PGA. Heatmaps showing the difference in mutant and wild-type embeddings generated for single mutation using sequence-only and joint sequence and structure embedding models, with regions having the highest difference in embedding values highlighted in blue and green. 3-D structures of considered proteins with the mutation site colored in red. The most-affected regions identified from the embedding space difference between wild-type and mutant are highlighted using the same color schemes.

**Supplementary Figure 6:**
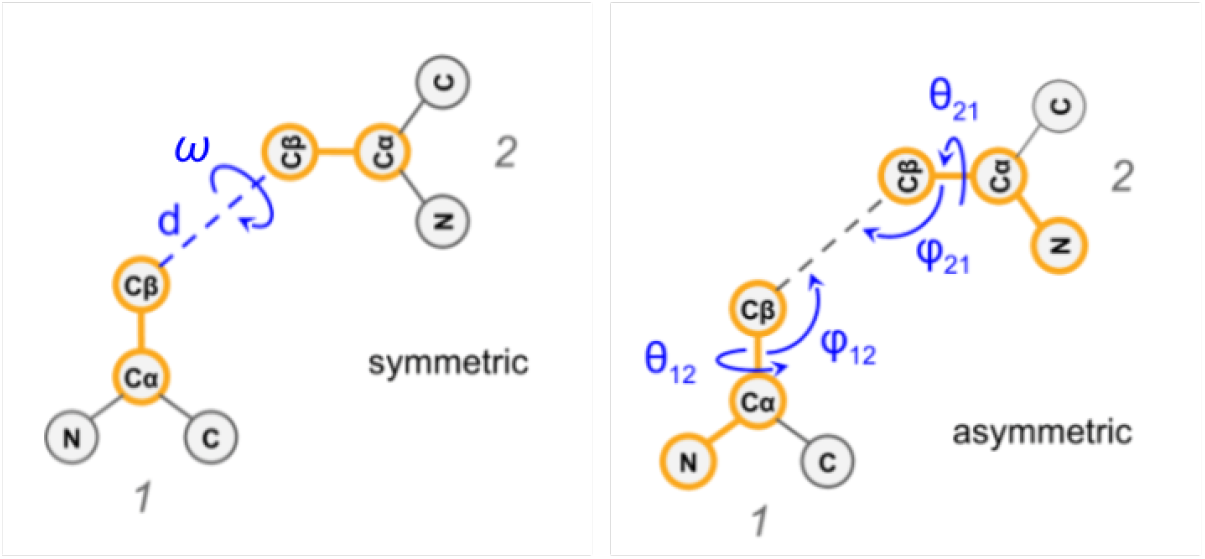
Inter-residue orientations used on edges for graph construction. Schematic depiction of inter-residue orientations (3 dihedral and 2 planar angles) used on edges connecting two residues in constructing the graph for input to SE(3) transformer (from Yang et al. (2020) [13]).

